# The unreasonable effectiveness of the total quasi-steady state approximation, and its limitations

**DOI:** 10.1101/2023.09.28.559798

**Authors:** Justin Eilertsen, Santiago Schnell, Sebastian Walcher

## Abstract

In this note, we discuss the range of parameters for which the total quasi-steady-state approximation of the Michaelis–Menten reaction mechanism holds validity. We challenge the prevalent notion that total quasi-steady-state approximation is “roughly valid” across all parameters, showing that its validity cannot be assumed, even roughly, across the entire parameter space — when the initial substrate concentration is high. On the positive side, we show that the linearized one-dimensional equation for total substrate is a valid approximation when the initial reduced substrate concentration *s*_0_*/K*_*M*_ is small. Moreover, we obtain a precise picture of the long-term time course of both substrate and complex.

## 1 Introduction

Enzyme assays serve multiple objectives. For catalytic reactions with established reaction mechanisms, these assays aim to assess the enzyme’s catalytic efficiency toward a substrate and to quantify the physical constants that define the catalytic process.

Within the realm of quantitative reaction analysis, unwavering rigor, precision, and accuracy in measuring these physical constants are essential. Achieving this necessitates the development of mathematical approximations for the rate laws that govern the reaction mechanisms. Furthermore, a profound comprehension of the experimental conditions that validate such approximations is indispensable [15].

The irreversible single-substrate, single-enzyme Michaelis–Menten reaction mechanism

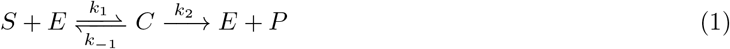

has been extensively explored. Numerous approximations for the rate law governing substrate depletion or product formation have been derived. On certain occasions, these approximations have been accompanied by stipulations that outline the conditions under which they are valid.

The ordinary differential equations that describe the time courses of the substrate, *s*, enzyme, *e*, and substrate-enzyme intermediate complex, *c*, concentrations are

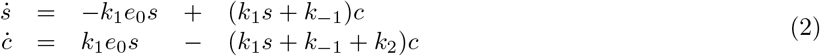

with parameters *e*_0_, *s*_0_, *k*_1_, *k*_−1_, *k*_2_. Here, *k*_1_, *k*_2_ and *k*_−1_ are rate constants, *s*_0_ is the initial substrate concentration, and *e*_0_ is the initial enzyme concentration. In the typical experimental assay, *c*(0) = 0, as there is no complex present at the beginning of the reaction.

The standard quasi-steady state assumption (sQSSA) for the Michaelis–Menten reaction mechanism assumes that the fast chemical species, the complex, is in quasi-steady state (QSS) with respect to the slow chemical species, the substrate. From *ċ* = 0, one derives the defining equation

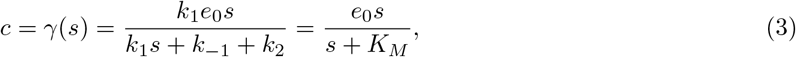

of the *c*-nullcline, and furthermore the familiar Michaelis–Menten equation

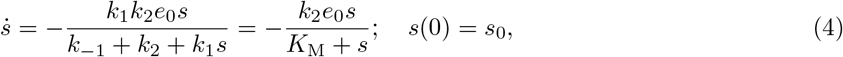

with the Michaelis constant K_M_ = (*k*_−1_ + *k*_2_)/*k*_1_. This reduction is based on the assumption of low initial reduced enzyme concentration, *e*_0_/K_*M*_ ≪ 1 (more precisely, *e*_0_ → 0 with K_*M*_ bounded away from 0) [5].

Moreover, in case of initial reduced substrate concentration *s*_0_/K_*M*_ being sufficiently small, one may further replace this by the linear approximation

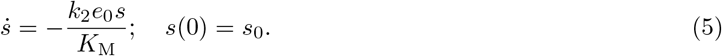

This linear approximation is of critical importance to estimate a fundamental kinetic parameter, *k*_2_/K_M_: the apparent second-order rate constant for the enzyme-catalyzed reaction. It is also known as specificity constant, because it determines the catalytic efficiency (or specificity) of the enzyme for each substrate. The higher the value of *k*_2_/K_M_ the more specific the enzyme is for a given substrate.

Recent work by Back et al. [1, Eq. (6)], Kim & Tyson [10], and Vu et al. [18, Eq. (8)] has introduced *linear tQSSA*. Maintaining the assumption of small *s*_0_/K_*M*_, but without restrictions on initial enzyme *e*_0_, they replaced substrate *s* by total substrate 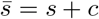, proposing to the reduced equation:

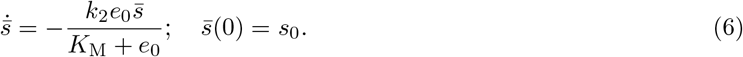

This “linear tQSSA” is grounded in the total quasi-steady-state assumption (tQSSA) first proposed by Borghans et al. [3]) and further developed by Tzafriri [17]. Equation (6) may be obtained from linearizing equation (9), which will be discussed below. It is important to note that aspects of their derivation rely on heuristical reasoning. In the course of the present paper we will derive equation (6) in a different way.

Eq. (6) can be expressed in terms of product concentration. By applying the conservation law 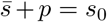, we arrive at:

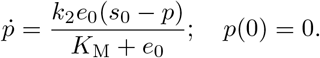

It is noteworthy that the specificity constant within the linearization (6), expressed as *k*_2_/(K_M_ + *e*_0_), diverges from the standard scenario by virtue of its dependence on the initial enzyme concentration.

The range of validity for the tQSSA is not clearly discernible in the existing literature. While Tzafriri [17] states it is “roughly valid” across all parameters, the precise meaning of this claim for the reduction remains unclear. Some interpretations treat this as a ”decent approximation” across the entire parameter space (e.g., Bersani and Dell’Acqua [2], who compare standard and total QSSA parameters). Kim and Tyson [10] take a more cautious approach, stating the tQSSA is “not too badly violated” for all parameters.

However, practical experience and numerical simulations suggest a broader applicability of the tQSSA. Notably, the work of Back et al. [1] and Vu et al. [18] provides strong support for the linear tQSSA, Eq. (6). This discrepancy between theoretical foundation and practical application compels us to echo E.P. Wigner’s famous phrase [19] and acknowledge the ”unreasonable effectiveness” of the tQSSA.

This manuscript aims to close this gap by providing a rigorous mathematical foundation for the linear tQSSA, Eq. (6), under the condition of low initial substrate concentration. Thus, a primary goal is to demonstrate that the effectiveness of the linear tQSSA is indeed reasonable.

Our analysis necessitates clarifying some misconceptions and limitations of the tQSSA. The claim of “rough validity” across all parameters, when interpreted as universal accuracy, is misleading. Additionally, the parameters proposed by Borghans et al. [3] and Tzafriri [17] do not accurately predict tQSSA validity.

Furthermore, the tQSSA’s heuristic derivation and associated criteria rely on an incorrect assumption of near-constancy of complex concentration. While this error does not negatively impact the long-term prediction of total substrate by Eq. (6), it leads to incorrect conclusions about the long-term behavior of free substrate and complex.

In contrast, our derivation of Eq. (6) is based on a mathematical approach (no heuristics) and only requires the assumption of low initial substrate (*s*_0_). This establishes a significantly broader range of applicability for the linear tQSSA. As an additional benefit, our derivation accurately predicts the time evolution of both free substrate and complex, leading to a better understanding of the tQSSA’s effectiveness.

An appendix in this paper offers a summary of findings regarding the efficacy of parameters proposed by Borghans et al. [3], and Tzafriri [17] for predicting the tQSSA validity, as analyzed through the lens of singular perturbation theory. While not essential for the main paper’s arguments and containing no groundbreaking results, this appendix may provide a helpful overview of scattered information from the literature.

## 2 Quasi-steady state approximation: Derivation and conditions

We start by reconsidering the conditions for the validity of the quasi-steady state approximations. These approximations are related to distinguished parameters, which guarantee (or are supposed to guarantee) their validity.

### 2.1 The standard Quasi-Steady State Approximation

For the case of the sQSSA, Segel and Slemrod [13] — following the earlier work of Briggs and Haldane — articulated the following argument: At low enzyme concentration *e*_0_, substrate depletion remains negligible during the short initial transient. Upon completion of this phase, the complex concentration will be nearly constant, implying ċ to be 0, approximately. Employing these conditions in conjunction with Eq. (2), they derived Eq. (4).

To assess the validity of this reduction, they employed partly heuristical timescale arguments to arrive at the dimensionless parameter:

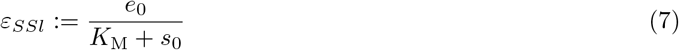

They posited that the sQSSA holds validity under the condition ε_*SSl*_ ≪ 1 (see, [13, Eq. (18a)]).

On this point, a brief digression regarding the notation “≪” is warranted. When we state “ε_*SSl*_ ≪1 implies validity of the sQSSA”, we mean the approximation error within the sQSSA falls below any specified bound, whenever ε_*SSl*_ is smaller than a corresponding constant determined by that bound.

However, it is crucial to note that this statement cannot be assigned a priori a fixed numerical interpretation (e.g., “ε_*SSl*_ ≤ 0.01”). Instead, we must address arbitrarily small bounds, necessitating the use of limit notation in mathematical terms—hence, we occasionally express this as the limit ε_*SSl*_→ 0. In the present paper, we therefore occasionally use the limit notation.

Another key point is that a statement such as “*e*_0_/(*s*_0_ +K_*M*_) →0” (or even more so, “*e*_0_/(*s*_0_ +K_*M*_) ≪1”) is equivocal, since there are substantially different ways to obtain this limit. For instance, one might have *s*_0_ and K_*M*_ constant, and let *e*_0_ → 0, or *e*_0_ and K_*M*_ constant and let *s*_0_ → ∞. It is important to remember that statements regarding limits, such as *e*_0_ →0, are independent of the chosen units.

Segel and Slemrod acknowledged this subtlety in their work. Their convergence proof [13, Section 6], which essentially demonstrates the existence of a singular perturbation reduction, was formulated under specific constraints: *s*_0_ must be bounded both below and above by positive constants, along with additional restrictions. Consequently, their work does not imply universal validity of the sQSSA under the condition ε_*SSl*_ ≪ 1, regardless of the manner in which this condition is achieved.

### 2.2 The total Quasi-Steady State Approximation

Extending and modifying the timescale arguments introduced in [13], Borghans et al. [3] assumed the total substrate 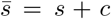 to be constant during a short transitory phase, and ċ ≈ 0 following that phase. Rewriting (2) for total substrate and complex and using ċ= 0, they obtained

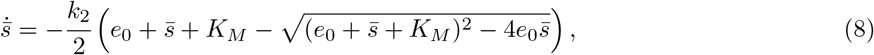

and — heuristically simplifying the right-hand side via Padé approximation — they arrived at

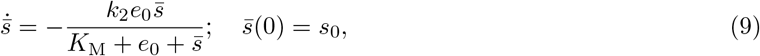

and the QSS relation

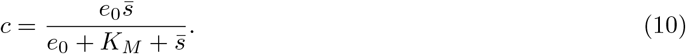

Considering (9) for low 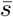 and keeping only the lowest order approximation on the right hand side, we obtain (6). Thus the linearized tQSSA may be seen as a consequence of tQSSA for low initial substrate.

In order to obtain conditions for validity of the approximation, Borghans et al. [3] established the dimensionless parameter

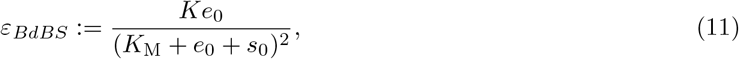

where K = *k*_2_/*k*_1_ is the Van Slyke-Cullen constant, and claimed that smallness of this parameter is necessary and sufficient for validity.

Tzafriri [17] took the same vantage point, but used different (more refined) arguments to obtain Eq. (9). He obtained the parameter

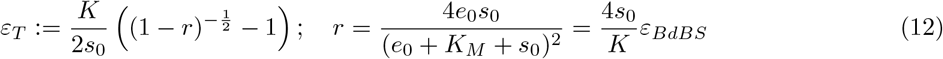

and claimed ε = ε_*T*_ ≪ 1 as a necessary and sufficient condition for validity [17, Eq. (32)]. One can verify that ε_*T*_ /ε_*BdBS*_ → 1 as ε_*BdBS*_ → 0, so ε_*T*_ ≪ 1 is equivalent to ε_*BdBS*_ ≪ 1. Based on the observation that ε_*T*_ is always bounded above by K/(4K_*M*_) 1/4, Tzafriri [17] states that tQSSA is always “roughly valid” for the Michaelis–Menten reaction mechanism; this is the origin of the notion. The following subsection will illustrate that “rough validity” — in the above sense — does not imply (even rough) approximate accuracy of the tQSSA.

From a mathematical perspective, we point out one gap in Tzafriri’s line of arguments (and also in the line of arguments in Borghans et al. [3]). In [17, equations (21) and (22)], Tzafriri claims with no further justification that ċ ≈ 0 for *all* times following the transitory phase. This claim was previously challenged by Schnell and Maini [12]. As will be seen below, we will show that it is incorrect to assume in the tQSSA that ċ ≈ 0 for *all* times following the transitory phase.

### 2.3 The system with high initial substrate

Borghans et al. [3] claim that ε_*BdBS*_ ≪ 1 is a necessary and sufficient condition for the validity of approximation (9), with Tzafriri [17] making a similar claim for (8) with ε_*T*_ ≪ 1. However, we demonstrate here that these assertions are inaccurate by examining the case of high initial substrate concentration (*s*_0_ → ∞).

It is evident that both ε_*BdBS*_ and ε_*T*_ approach zero as *s*_0_ → ∞, while assuming other parameters remain fixed at positive values. Adhering to their criteria, this would imply the validity of tQSSA in such scenarios.

Yet, we present numerical simulations with high *s*_0_ that reveal shortcomings in the tQSSA approximation. **Example**. We choose parameters (in arbitrary units): *s*_0_ = 10^4^, *e*_0_ = 10, *k*_1_ = 0.5, *k*_2_ = 5.0, and *k*_−1_ = 0.1. So we have K = 10, K_*M*_ = 10.2 and ε_*BdBS*_ ≈ ε_*T*_ ≈ 10^−6^. So, both parameters predict good approximation of the solution obtained from the full system (2) by the solution of (9), or (8). But this is not the case as illustrated in Figures 1 and 4 for the time courses, and Figures 2 and 5 for the ratios of the approximations. One may obtain a misleading first impression by looking only at the situation when 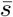 is still high, and the approximation seems to be good. However, the discrepancy becomes clearly notable when 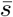 decreases, and becomes pronounced in the interesting range 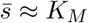 K_*M*_. (The discrepancy between solutions of (9) and (8) is due to the approximation error of the linearization.) In Figure 3, we compare the time course of complex concentration, and the ratio of complex and total substrate concentrations, obtained from systems (2), and (9).

**Figure 1:**
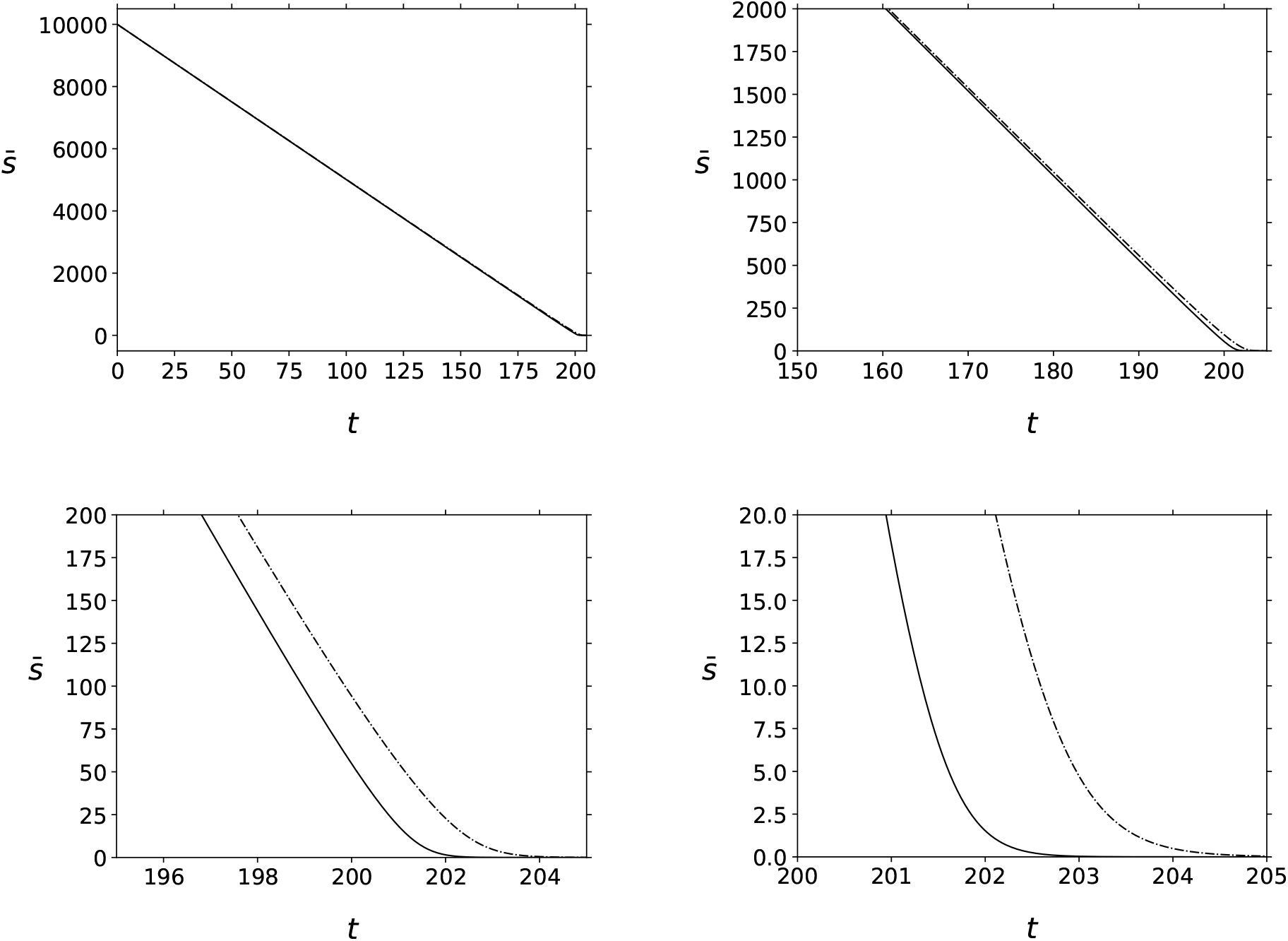
Large *s*_0_ does not enhance the accuracy of (9). In this simulation, the solid black curve is the time course of 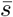 obtained by numerically integrating the mass action equations (2). The dashed/dotted curve is the time course of 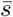 obtained via the numerical integration of (9). While the approximation (9) is good for large 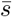, it becomes less reliable as 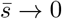.

**Figure 2:**
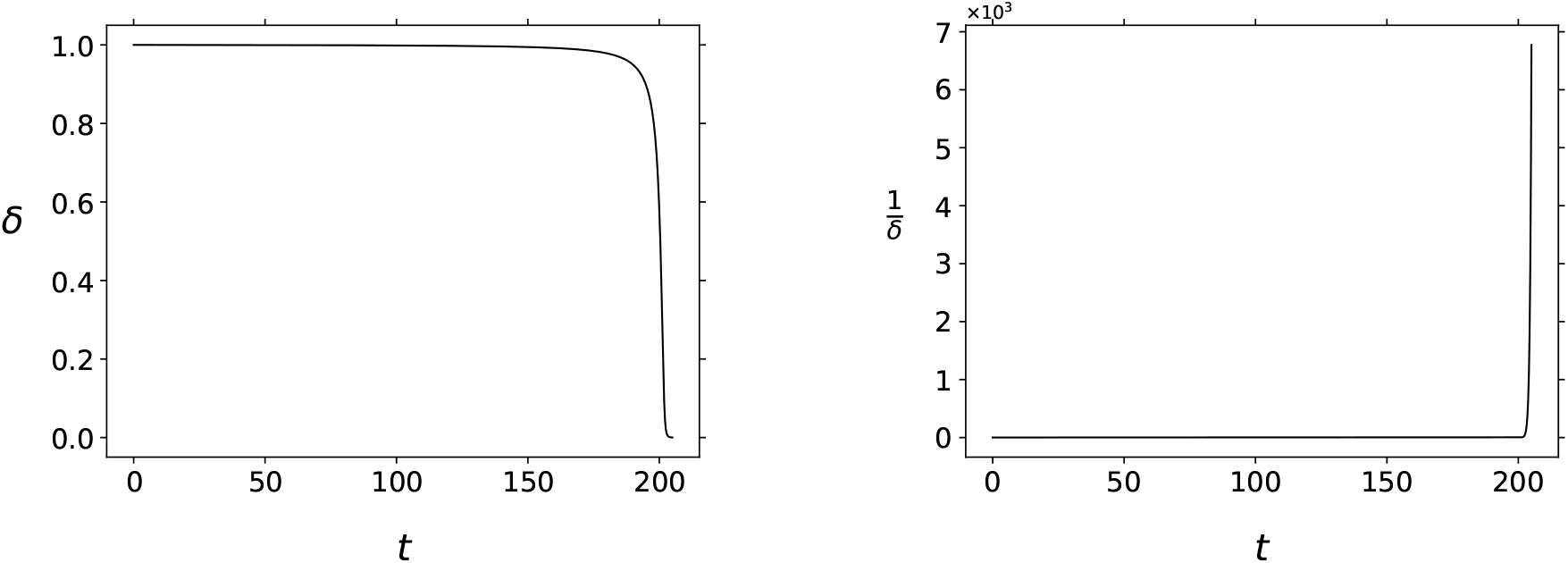
Large *s*_0_ does not enhance the accuracy of (9) and (10). “δ” denotes the ratio of 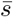 obtained from the mass action equations to 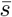 obtained via integration of (9).

**Figure 3:**
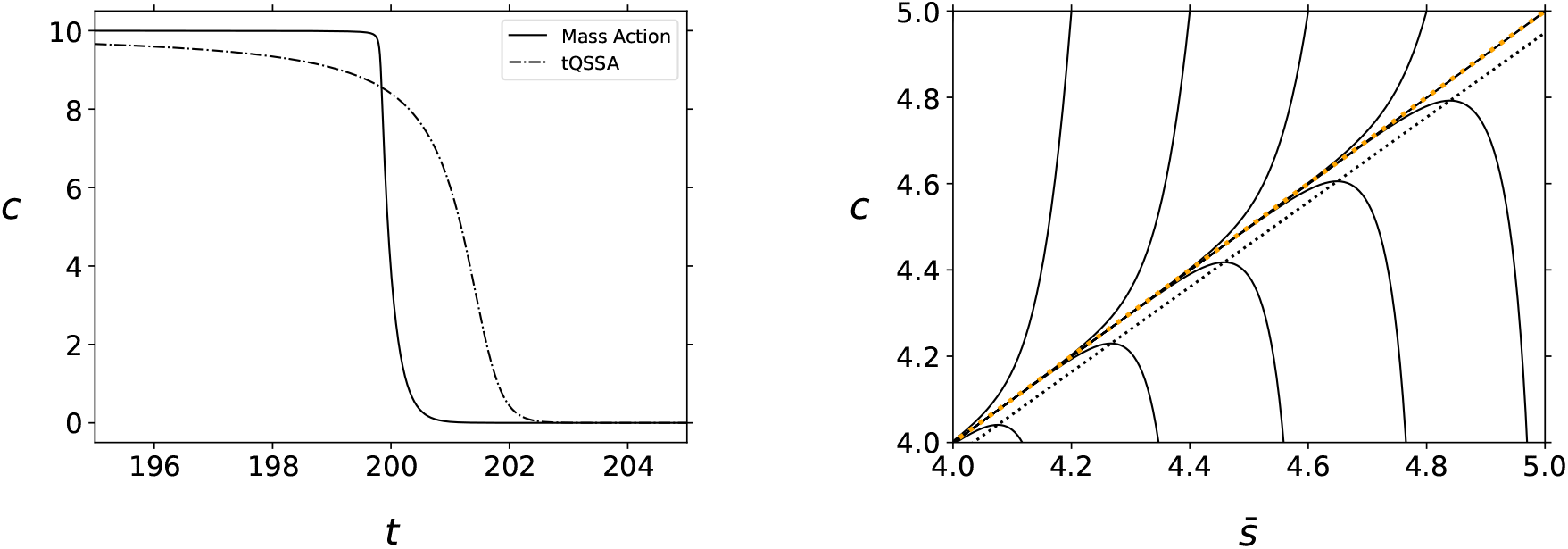
Large *s*_0_ does not enhance the accuracy of (9) and (10). In both panels, the solid black curve is the time course of complex concentration, *c*, obtained via numerical integration of the mass action equations (2). The dashed/dotted curve is the time course of *c* given by (9)–(10). Clearly, the differential– algebraic system, (9)–(10), eventually fails to approximate the time course of 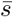 and *c*. In this simulation, various trajectories (obtained from the numerical simulation of the mass action system (2)) are plotted in the phase plane. Initial conditions for 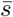 vary from 4.0 ≤ *s*_0_ ≤5.0 and *s*_0_ = 10^4^. The dotted black line is the *c*-nullcline; the dotted orange line corresponds to *s* = 0.

**Figure 4:**
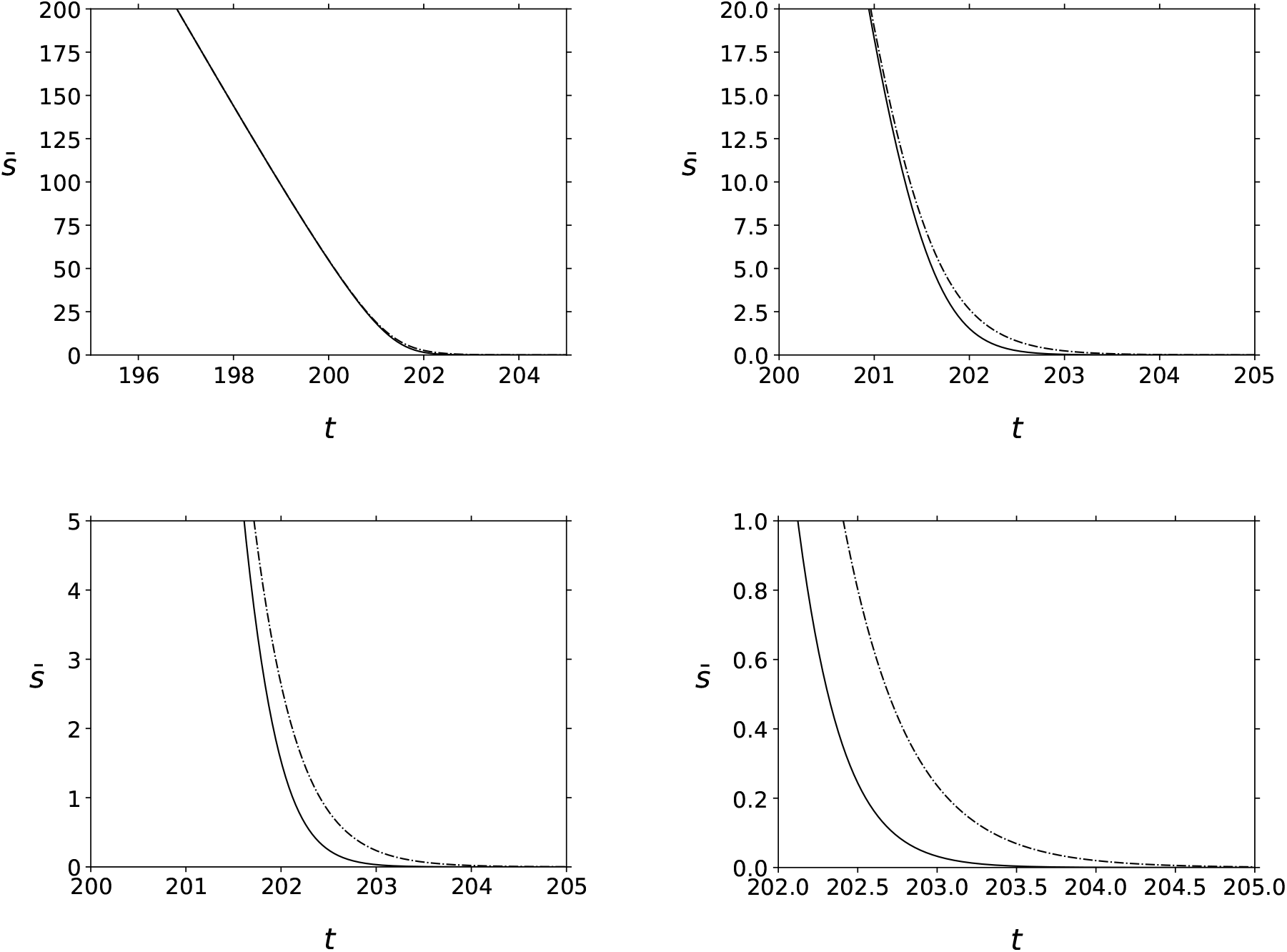
Large *s*_0_ does not enhance the accuracy of (8). In this simulation, the solid black curve is the time course of 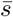 obtained by numerically integrating the mass action equations (2). The dashed/dotted curve is the time course of 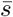 obtained via the numerical integration of (8). While the approximation (8) is good for large 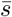, it becomes unreliable as 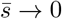.

**Figure 5:**
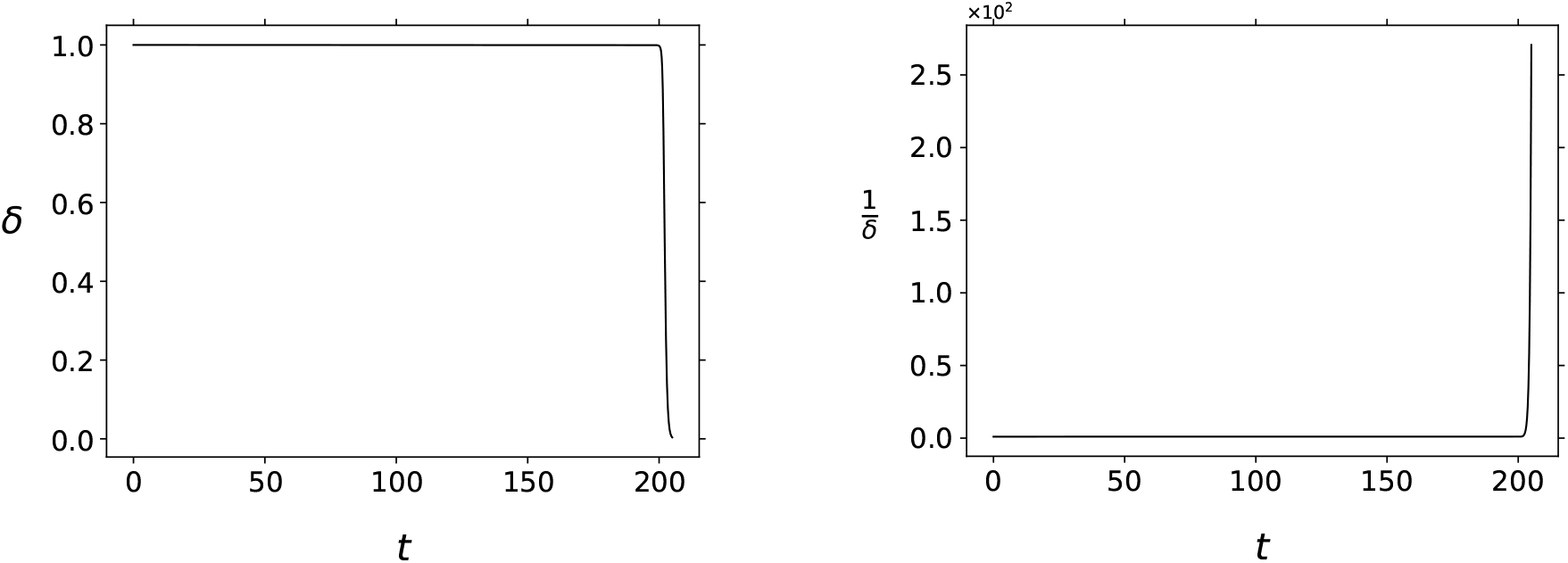
Large *s*_0_ does not enhance the accuracy of (8). “δ” denotes the ratio of 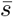 obtained from the mass action equations (2) to 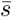 obtained via integration of (8).

We draw some conclusions from this example:

1. The parameters ε_*BdBS*_ and ε_*T*_ do not correctly predict validity of tQSSA.
2. The statement that the tQSSA is “roughly valid” for all parameters cannot be interpreted as — even rough — accuracy of the approximation. And one sees more: Tzafri’s claim [17, end of subsection 2.1] that tQSSA is a good approximation when ε_*T*_ ≤ 0.1 is falsified by the example, where ε_*T*_ ≈ 10^−6^.
3. From Figure 3, one sees that trajectories approach the origin tangent to the line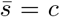, thus *s*= 0, independent of initial substrate, even for *s*_0_ = 10^4^. Hence, large *s*_0_ does not ensure that phase plane trajectories stay close to the *c*-nullcline for all time.

## 3 The linearization of the reaction dynamics and tQSSA

Our discussion and simulations reveal that tQSSA generally is not generally valid at high initial substrate concentrations. This finding extends to moderately high *s*_0_ through further simulations. Consequently, from a mathematical perspective, focusing on the case of low *s*_0_ becomes natural. Furthermore, the assumption of low initial substrate forms the cornerstone of Back et al. [1] and Vu et al. [18], whose work focuses on pharmacokinetic applications. In the case of *s*_0_/K_*M*_ ≪ 1, both ε_*BdBS*_ and ε_*T*_ tend to the parameter

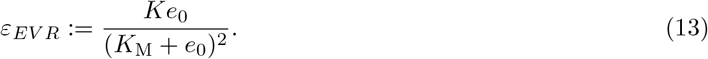

Approximately four times this value, 4ε_*EV R*_, serves as an approximation for the eigenvalue ratio of the linearization at the stationary point (see, for details, [5, Proposition 1 and Remark 2]). In the example discussed in the previous section, we find ε_*EV R*_ ≈1/4, indicating an eigenvalue ratio close to 1. Additionally, the right-hand sides of equations (8) and (9) tend towards the right-hand side of Eq. (6) near the stationary point 0.

We will show below that small ε_*EV R*_ leads to an approximation of (2) by a two-dimensional linear system, from which (6) can be obtained. Building upon the findings of [5], one can argue that while not universally applicable, tQSSA holds validity under conditions of low initial reduced substrate concentration. This characteristic is crucial in explaining the seemingly “unreasonable effectiveness” of tQSSA. Consequently, ε_*EV R*_ should be seen as the relevant parameter, rather than ε_*BdBS*_ or ε_*T*_.

### 3.1 A direct derivation of the linear tQSSA

Consider the linearization of the governing equation for the Michaelis–Menten reaction mechanism (1) at the stationary point 0

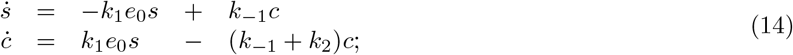

thus the right-hand side is replaced by the linear Taylor approximation at the stationary point 0. As is known from the theory of ordinary differential equations (see e.g. Sternberg [14, Theorem 2], and also Chicone [4, Theorems 4.1, 4.8 and 4.14]), near the stationary point the systems (2) and (14) are smoothly equivalent, hence replacing solutions of the former are by solutions of the latter provides a valid local approximation. The eigenvalues of the matrix in (14) are

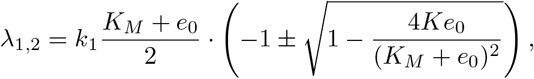

with λ_1_ of smaller absolute value. With corresponding eigenvectors v_1,2_, every solution of the differential equation has the form

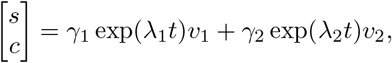

with constants γ_1,2_ determined by the initial values. Now when |λ_1_/λ_2_ | ≪1 (equivalently ε_*EV R*_ ≪ 1), then the second term in the above sum will decay quickly, thus after a short initial phase we have a good approximation by

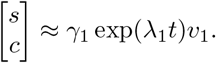

Loosely speaking, the dominating timescale for the degradation of any species (regardless of the initial conditions) is |λ_1_| ^−1^, given small ε_*EV R*_.

In turn, by differentiation one obtains

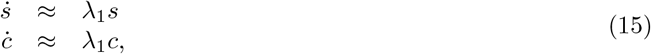

and furthermore with 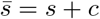 we arrive at

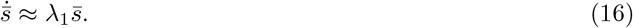

Finally, given small ε_*EV R*_, one has

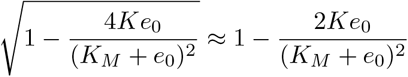

by standard Taylor approximation, which implies

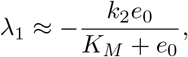

hence

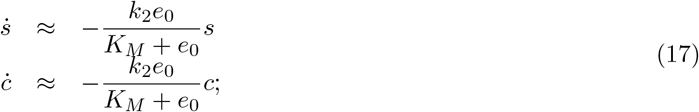

in particular (16) agrees with (6). Thus, the reduction of the Michaelis–Menten equation to (6) near 0 is justified by a general property of differential equations and their linearizations at stationary points.

### 3.2 Comparing the tQSSA derivations

While Borghans et al. [3] and Tzafriri [17] derived Eq. (6) from the tQSSA in a mathematically problematic way, it nevertheless yielded a correct reduction. This might tempt some readers to conclude that the derivation’s flaws do not matter. Indeed, it is logically possible that incorrect arguments lead to correct conclusions. However, we now point out that the same incorrect derivation also leads to an incorrect conclusion here.

We discuss the case of small *s*_0_ and small ε_*EV R*_ in greater detail, looking explicitly at the eigenvector *v*_1_ for λ_1_. By a straightforward calculation, the direction of the slow eigenspace is given by

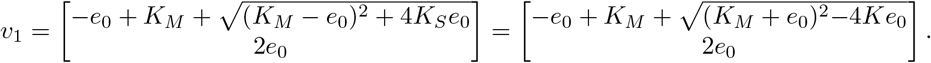

Moreover, given ε_*EV R*_ ≪ 1, we have the approximation by Taylor expansion (as above)

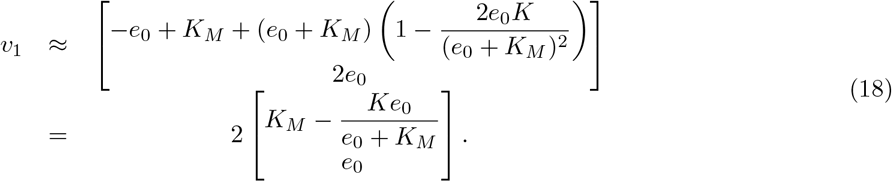

This shows that ċ ≈ 0 in the long-time regime only if *e*_0_ → 0, which is the sQSS case. Otherwise, the ratio of substrate and complex is asymptotic to

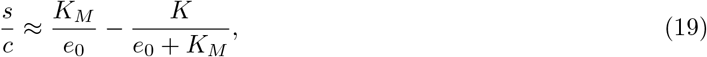

hence finite. This contradicts the claim of both Borghans et al. [3] and Tzafriri [17] that ċ ≈ 0 in the whole time period following a short transitory phase, and which is a cornerstone of their arguments. (As noted before, this assumption is a critical problem in their arguments.) Stated from a different perspective, 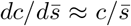 in the linear tQSSA after the transient, so the statement ċ ≈ 0 only applies when *c ≪ s* in the approach to the stationary point at the origin.

We proceed to compare the above to the usual derivation of the tQSSA reduction. In 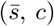 coordinates we have

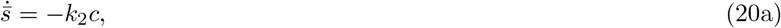

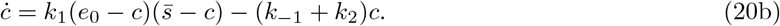

The tQSSA is formally given by

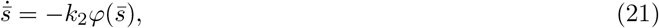

where 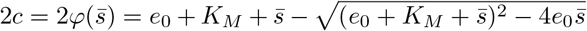 describes the *c*-nullcline in the 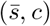 coordinate system. In this coordinate system,

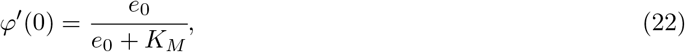

and so near the stationary point, the linearized tQSSA yields

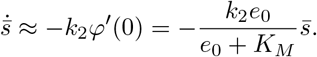

This is always in agreement with the linearization of the full system in the presence of timescale separation. However, in (*s, c*) coordinates the *c*-nullcline is given by (3), and with the quasi-steady state assumption for complex, the solution trajectory should be close to this nullcline near the stationary point 0. In particular, for the tangent at 0 one has

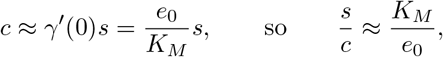

which is markedly different from (19) whenever *e*_0_ is not small. So, while providing the correct timescale, the approach in [10] generally provides an incorrect asymptotic ratio of substrate and complex as illustrated in Figure 6.

**Figure 6:**
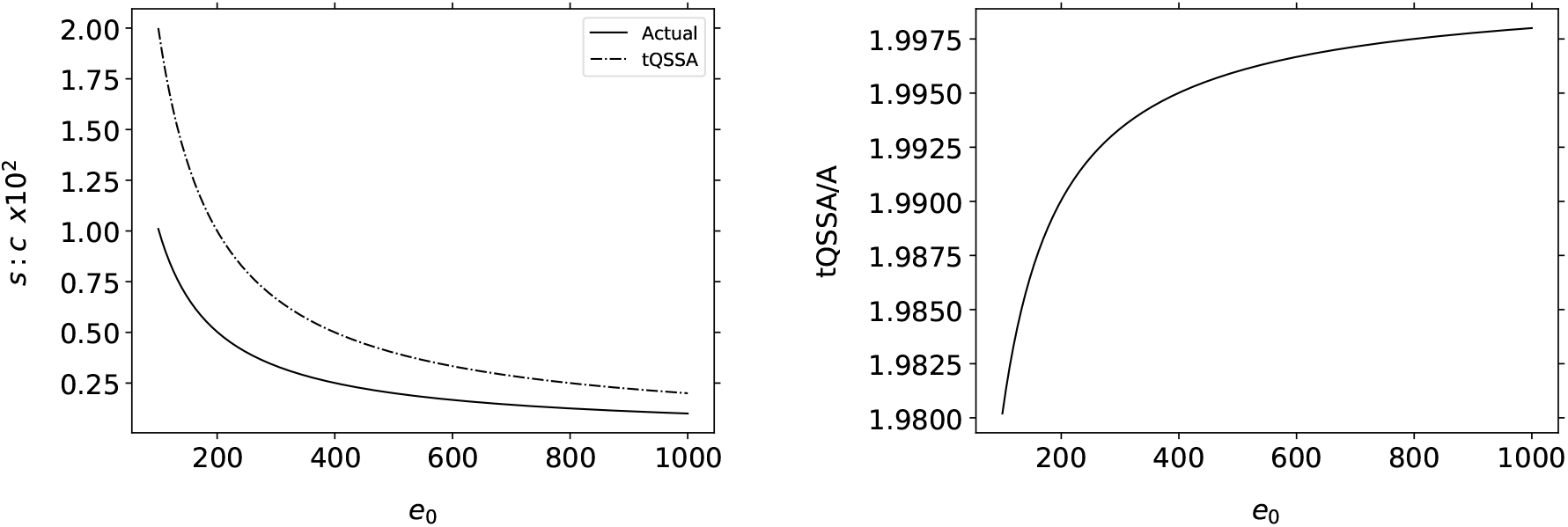
The linear tQSSA can fail provide the correct asymptotic ratio of substrate to complex. A frequently reported advantage of the tQSSA is its purported accuracy at high enzyme concentration. However, *accuracy* in this context is limited to the approximation of the slow timescale. The tQSSA tends to overestimate the asymptotic ratio of substrate to complex. The parameters (in arbitrary units) utilized in this figure are: *k*_1_ = *k*_2_ = *k*_−1_ = 1.0 and *e*_0_ is varied from 100 to 1000. Left: The solid black curve is the actual asymptotic ratio of substrate to complex, *s* : *c*, obtained from integration of (2). The dashed/dotted curve is the asymptotic ratio of substrate to complex provided by the linear tQSSA (6). Right: The black curve is *s* : *c* obtained from the linear tQSSA, divided by *s* : *c* obtained from the slow eigenspace as a function of *e*_0_. As *e*_0_ increases, we see that the tQSSA overestimates *s* : *c* by a factor of roughly 2.

What exactly is positive about the correct linearization approach? Linearization of (9) offers a crucial advantage: it provides more (and correct) information about the system’s time evolution.

## 4 Discussion

In the mathematical enzymology community, it is generally accepted that the tQSSA is “roughly valid” [17], but the actual meaning of this statement is not very clear. The validity of the tQSSA has been explored through heuristic approaches and numerical simulations. It appears that Vu et al. [18] in their recent work do not explicitly address the necessity of small ε_*EV R*_ for the validity of the tQSSA. They might be presuming that this condition is universally fulfilled, or perhaps they noted by inspection of [5, especially Figure 2] that the condition holds true for most of the parameters (in a well-defined sense).

In this note, we provide a solid foundation for the tQSSA approximation (6), and show its validity when *s*_0_/K_*M*_ ≪ 1. It is easy to see that there is no contradiction between the tQSSA approximation (6) and the sQSSA approximation (5) as (6) tends (5) for *e*_0_→0. However, the tQSSA predicts a specificity constant dependent on the initial enzyme concentration at high enzyme concentrations. This noteworthy aspect has not yet been observed in experimental settings.

Two mathematical issues remain unresolved. First, the linearization theorem for differential equations only holds locally near stationary points, offering no information about its range validity. A forthcoming paper, using advanced mathematical tools, will demonstrate that the only limitation requires low initial substrate concentrations; no further restrictions exist. Second, for linear tQSSA, we need to determine starting values for both *s* and for *c*, or for 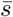 at the beginning of the slow regime, following the initial fast transient. This, too, will be addressed in the forthcoming paper.

## Acknowledgments

We thank four anonymous reviewers for critically reading the manuscript, and for valuable comments. SS would like to thank the Isaac Newton Institute for Mathematical Sciences for support and hospitality during the programmed *Collective Behaviour – MMVW02* when work on this paper was partially undertaken. As a result, this paper was partially supported by EPSRC Grant Number EP/R014604/1.

## 5 Appendix: On the parameter of Borghans et al. [3]

The tQSSA parameter (11) proposed in [3], which can be rewritten as

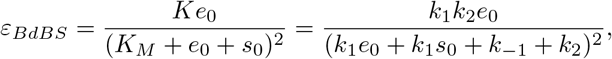

is widely accepted in the mathematical biology community, as a predictor for validity of the reduction (9). While we have seen above that the prediction fails when *s*_0_ → ∞, many results in the literature support the efficacy of the condition ε_*BdBS*_ ≪ 1 in other scenarios. This may be seen as another aspect of the “unreasonable effectiveness” of the tQSSA.

In this appendix we will discuss all parameter combinations which yield ε_*BdBS*_ ≪1, and compare the tQSSA to reductions whose validity is guaranteed by singular perturbation theory. As noted earlier the condition ε_*T*_ ≪ 1 due to Tzafriri [17] is equivalent to ε_*BdBS*_ ≪1; so it suffices to discuss one of them. While these considerations are not central to the present paper, a complete listing may be useful for practitioners. We will not carry out all the mathematical details, but rather rely on, or report, facts and procedures known from the literature.

To start, one needs to break down the condition ε_*BdBS*_ ≪ 1 into manageable bits. We interpret this condition as ε_*BdBS*_ → 0, thus a condition on a limit, which needs to be taken in a well-defined manner. Now ε_*BdBS*_ →0 is possible only if the numerator (thus one of the factors in the numerator) approaches zero, or if one of the terms in the denominator approaches infinity. Taking limits we will keep all parameters except one fixed at positive values, with only the remaining one approaching zero or infinity. One may verify that more complicated scenarios lead to similar approximations, or to degenerate ones with 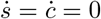. We consider all possible nondegenerate cases, even if some may be of little practical interest:

i. In the case when all concentrations and reaction rates remain bounded, one sees that ε_*BdBS*_ → 0 only if *k*_1_→ 0, or *e*_0_ →0, or *k*_2_→0. These cases are well known and lead to singular perturbation reductions (see Goeke et al. [8, 9]), which are certainly valid by results of Tikhonov [16] and Fenichel [6]. One verifies that — up to higher order terms in the respective small parameter — the singular perturbation reduction agrees with (8), and furthermore with (9) in the case *e*_0_ → 0 (and also when *k*_1_ → 0).
ii. We have seen in subsection 2.3 above that tQSSA fails when *s*_0_ → ∞.
iii. The case *e*_0_ → ∞ leads to ε_*BdBS*_ → 0, and it is presented in detail via numerical studies in Borghans et al. [3, Section 3] with the conclusion that tQSSA is valid for high *e*_0_. We discuss the case of high *e*_0_, from the singular perturbation perspective, following Noethen and Walcher [11, Example 3.2]. Take 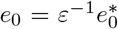, where ε is a dimensionless parameter and 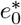 is understood to be of unit magnitude. The mass action equations for *s* and *c* in perturbation form are

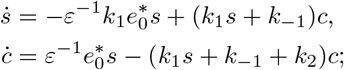

which is a slow-fast system with natural time being the slow time. This system is amenable to a singular perturbation reduction; see [7] for details. With fast time T = ε^−1^t, we have the layer problem in the form

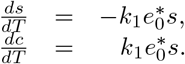 Thus, the critical manifold is simply the line *s* = 0, and the QSS species here is *s* (with 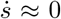 0 in the slow regime), not *c* as assumed in [3] and [17]. By the computational procedure from [7, Theorem 1, Eq. (8)] the slow dynamics is given by

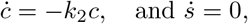

which in turn implies 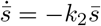. This is also recoverable from (8) in the limiting case *e*_0_ → ∞, and also from (15), since λ_1_ → −*k*_2_ as *e*_0_ → ∞. So, the tQSS reduction provides a correct timescale.
iv. Next, we discuss the case *k*_1_ → ∞ leading to ε_*BdBS*_ → 0. The tQSS reduction (9) yields

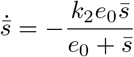

in the limit *k*_1_→∞. From the singular perturbation perspective, we take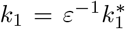, where ε is a dimensionless parameter and 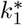 is understood to be of unit magnitude. The mass action equations for *s* and *c* in perturbation form are

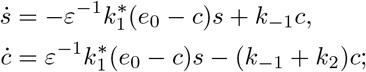

which again is a slow-fast system with natural time being the slow time. Here the critical “manifold” is given by (*e*_0_ − *c*)*s* = 0, hence is the union of the lines *c* = *e*_0_ and *s* = 0. This yields two singular perturbation reductions. Near the line *c* = *e*_0_, one gets

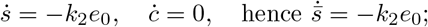

which clearly can hold only for a limited time. Near *s* = 0 one finds

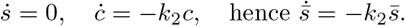 We see that the tQSSA provides an incorrect reduction here. The case *k*_−1_→ ∞ yields 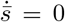, and likewise the singular perturbation reduction yields a reduced equation with trivial right-hand side. Thus, both reduction methods agree, but yield a degenerate case which would require further analysis.
v. Finally, we discuss the case *k*_2_ → ∞ leading to ε_*BdBS*_ → 0. The tQSSA with (9) yields

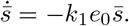 From the singular perturbation perspective, let 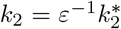, where ε is a dimensionless parameter and 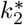 is understood to be of unit magnitude. The mass action equations for *s* and *c* in perturbation form are

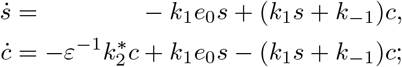

again a slow-fast system with natural time being the slow time. Here the critical manifold is given by *c* = 0; so *c* is the QSS variable. The reduction procedure yields

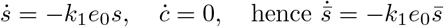

in agreement with the tQSSA.

**Table 1:**
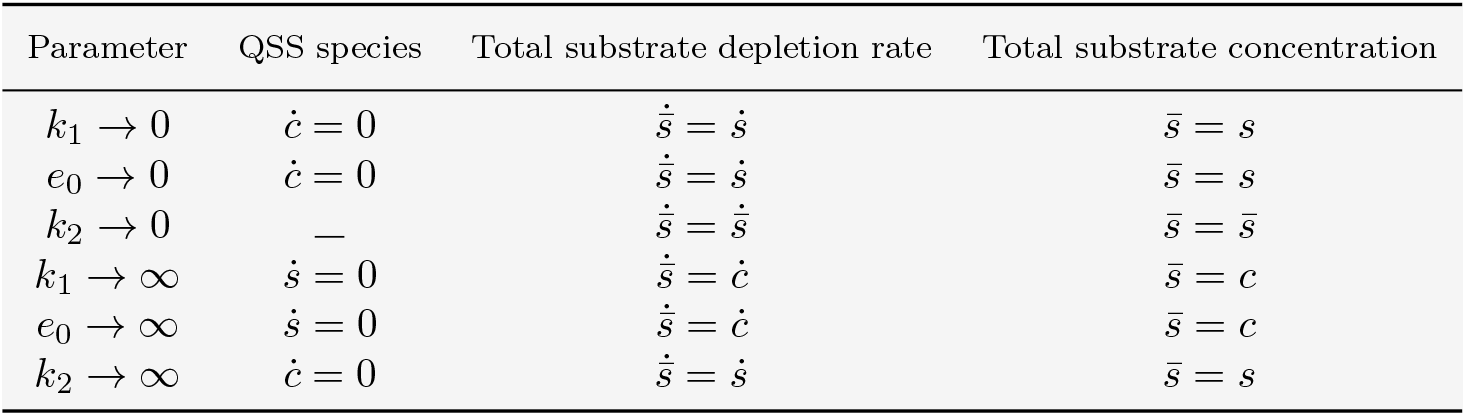
Quasi-steady-state behavior: *Small* and *large* parameters.

Table 1 provides an overview of our results, and some additional information about the reductions. Briefly one could say that the parameter ε_*BdBS*_, which was obtained from in part heuristic reasoning, does not correctly predict the reduction in all possible cases, and an independent verification (or falsification) — as outlined above — is necessary. But the parameter provides correct predictions for a number of practically relevant cases.

